# Metagenomic sequencing provides insights into the location of microbial detoxification in the gut of a small mammalian herbivore

**DOI:** 10.1101/299198

**Authors:** Kevin D. Kohl, Kelly F. Oakeson, Teri J. Orr, Aaron W. Miller, Jennifer Sorensen Forbey, Caleb D. Phillips, Colin Dale, Robert B. Weiss, M. Denise Dearing

## Abstract

Microbial detoxification of plant defense compounds influences the use of certain plants as food sources by herbivores. The location of microbial detoxification along the gut could have profound influences on the distribution, metabolism, and tolerance to toxic compounds. Stephen’s woodrats (*Neotoma stephensi*) specialize on juniper, which is heavily defended by numerous defensive compounds, such as oxalate, phenolics, and monoterpenes. Woodrats maintain two gut chambers harboring dense microbial communities: a foregut chamber proximal to the major site of toxin absorption, and a cecal chamber in their hindgut. We performed several experiments to investigate the location of microbial detoxification in the woodrat gut. First, we measured levels of toxins across gut chambers. Compared to food material, oxalate concentrations were immediately lower in the foregut chamber, while concentrations of terpenes remain high in the foregut, and are lowest in the cecal chamber. We also conducted metagenomic sequencing of the foregut and cecal chambers to compare microbial functions. We found that the majority of genes associated with detoxification functions were more abundant in the cecal chamber. However, some genes associated with degradation of oxalate and phenolic compounds were more abundant in the foregut. Thus, it seems that microbial detoxification may take place in various chambers depending on the class of chemical compound. We hypothesize that the location of microbial detoxification could impact the tolerance of animals to these compounds, which may have ecological and evolutionary consequences.

## INTRODUCTION

Mammalian herbivores are often challenged by the consumption of toxic plant secondary compounds (PSCs) that negatively impact their physiology in various ways [1]. In response, mammalian herbivores have evolved myriad adaptations to cope with dietary toxins, such as altered behavior, efflux pumps in the gut, and enhanced liver detoxification [1]. Recently, we demonstrated that the gut microbiota of herbivores also play an important role in allowing them to consume PSCs [2]. However, it is still unclear where the process of microbial detoxification occurs in the gut.

It has been proposed that microbial detoxification should occur early in the gastrointestinal tract, to facilitate rapid degradation of PSCs and minimize the potential for their absorption in the digestive tract. In fact, it has been hypothesized that the rumen evolved first for detoxification and was later adopted for processes of fermentation [3, 4]. Examples of animals in which foregut detoxification has been documented for particular PSCs include reindeer (*Rangifer tarandus;* [5] that have rumen communities that degrade usnic acid, a toxin produced by lichen, and hoatzins (*Opisthocomus hoazin*), that maintain communities of microbes in their foregut capable of metabolizing saponins [6]. There is also evidence of microbial detoxification processes occurring in the hindguts of some animals. For example, Sage-grouse (*Centrocercus urophasianus*), specialize on toxic sagebrush (*Artemsia* spp.) and maintain a cecal microbiota enriched in detoxification genes [7]. Additionally, the hindguts of koalas (*Phascolarctos cinereus*) harbor microbes capable of degrading phenolic PSCs [8]. Understanding the location of microbial processes in the gut is likely to be important in the context of animal ecology. For example, the location of fermentation (foregut vs hindgut), seems to facilitate or constrain evolution of herbivore body size [9] and dietary niche breadth [10]. The location of microbial detoxification could have similar impacts on animal ecology.

In contrast to many above described examples, woodrats (genus *Neotoma*) are particularly interesting from a digestive standpoint as these herbivorous rodents possess both a foregut chamber and a large cecal chamber in their hindgut. Both the foregut and cecal chambers harbor dense communities of microbes and high concentrations of volatile fatty acids [11], which are the microbial products of fermentation. Following the hypothesis that the bovine rumen first evolved for detoxification regarding the rumen [3, 4], we predict that microbial detoxification of PSCs might occur prior to the small intestine in the woodrat gut such that the animal experiences limited absorption and systemic exposure to toxic compounds.

To evaluate this hypothesis, we conducted a series of experiments focusing on a species of woodrat with a highly specialized diet. Stephen’s woodrat (*Neotoma stephensi*) is a dietary specialist, with juniper (*Juniperus monosperma*) composing ~90% of its diet [12]. Juniper defends itself from herbivory with various toxic compounds, including monoterpenes [13], phenolics [14, 15], and oxalate [16]. We have demonstrated that microbes aid in the detoxification of phenolics [2] and oxalate in other woodrat species [17, 18]. For example, some populations of the white-throated woodrat (*N. albigula*) specialize on cactus (*Opuntia* spp.), which is rich in oxalate, but lacking in other defensive compounds. The foregut microbial communities of these *N. albigula* contain several species of oxalate-degrading microbes [19]. There has also been metagenomic evidence obtained from other systems suggesting that microbes aid in the detoxification of terpenes (mountain pine beetles: [20]; sage grouse: [7], as well as experimental work demonstrating the detoxification function of the gut microbiota of pine weevils [21]). However, the detoxification contribution of the microbiome has not been compared between the foregut and hindgut in any wild vertebrate herbivore.

We addressed several questions within the woodrat-juniper system. First, we measured concentrations of two classes of toxins (oxalate and monoterpenes) across regions of the gut. We predicted that if detoxification is occurring proximally in the gut, then toxins concentrations in the foregut would be significantly lower compared to food material. Next, we conducted a series of metagenomic analyses to inventory and compare the functions of microbial communities. We first compared the metagenomes of foregut communities between the juniper specialist *N. stephensi* and the cactus specialist *N. albigula* and predicted that if the foregut microbiota of *N. stephensi* were specialized for detoxification of juniper, which contains oxalate, monoterpenes, and phenolics, we would observe enrichment of detoxification genes compared to the foregut of *N. albigula* that is exposed primarily to the oxalate in cactus. We also compared the metagenomes of foregut and cecal chambers within *N. stephensi* and predicted that the metagenome of the foregut chamber would be enriched in detoxification genes compared to the cecal communities.

## MATERIALS AND METHODS

All experiments below were approved by the University of Utah Institutional Animal Care and Use Committee (IACUC) under protocols #12-12010 and #16-02011.

### Toxin Levels Throughout the Gut

We conducted a feeding trial on *N. stephensi* to document changes in toxin concentrations across gut regions. Six individuals of *N. stephensi* were fed increasing concentrations of *J. monosperma* (3 days on 25%, 3 days on 50%, 4 days on 75%). On the last morning, animals were euthanized using isoflurane. Contents of each gut region (foregut, stomach, small intestine, cecum, large intestine, feces) were collected. A small portion of wet gut contents were placed in airtight headspace autosampler vials and frozen at −20°C until analysis. Concentrations of individual monoterpenes were determined using headspace gas chromatography with an Agilent 7694 headspace sampler coupled with an Agilent 6890N gas chromatograph (GC). One mL of headspace gas was injected into a J&W DB-5 capillary column (30 m × 250 μm × 0.25 μm). Operating conditions for the headspace sampler were: oven temperature at 100°C, loop temperature at 110°C, transfer line temperature at 120°C, a vial equilibrium time of 20 min, a pressurization time of 0.20 min, a loop fill time of 0.50 min, a loop equilibrium time of 0.20 min, and an injection time of 0.50 min. Operating conditions for the GC were: splitless injector at 250°C, flame ionization detector at 300°C, oven temperature at 40°C for 2 min, then increasing 3°C/min to 60°C, then increasing 5°C/min to 120°C, then increasing 20°C/min to 300°C, and held at 300°C for 7 min. The make-up gas was nitrogen and the carrier gas was helium. The inlet pressure was 80 kPa with a flow rate of 1.0 mL/min. Retention times and peak areas of individual monoterpenes were calculated using Agilent OpenLab version A.01.05. We verified the relative retention times of individual monoterpenes using standards, and estimated the relative concentrations of individual monoterpenes in the intestinal content material by comparing AUC of peaks in diet or gut samples to AUC of a known volume of 10 mg/mL monoterpene standards.

Remaining gut contents were placed in weighing boats and dried at 40°C for two nights. The dried gut contents were then used to measure oxalate concentrations using a previously established protocol [22]. Roughly 0.4g of dried gut contents were ground and added to 5 mL of **6** N H_2_SO_4_ to solubilize the oxalate. After 15 min, 25 ml of distilled water was added, and the entire solution was filtered through grade 4 Whatman filter paper. The filtrate was brought up to a pH of 7 with NaOH, and 0.1 g of CaCl_2_ was added to precipitate the oxalate. The samples were centrifuged and decanted. After centrifugation, a volume of distilled water equal to that recovered after filtration was added, and the samples were titrated.

### Sample collection for metagenomics

We compared the metagenomic sequencing results across species and gut regions to investigate differential abundance of microbial functions. We collected individuals of the cactus specialist *N. albigula*, from Castle Valley, UT, USA (38°30’ N, 109°18’ W). Individuals of *N. stephensi* (different individual from those described in the experiment above) were collected from near Wupatki National Monument, AZ, USA (35°30’ N, 111°27’ W). Animals were immediately transported back to the University of Utah animal facility, provided with native diets: either cactus pads or juniper overnight (for *N. albigula* and *N. stephensi*, respectively), and euthanized and dissected on the following morning. Contents of the foregut chamber were collected from three individuals of both species, and the contents of the cecal chamber were collected from *N. stephensi*. We sequenced the metagenome of each sample (described below), and conducted several comparisons. First, we compared the foregut metagenomes of *N. albigula* and *N. stephensi*, with the prediction that the foregut of *N. stephensi* would be enriched in genes associated with detoxification of juniper, which contains more chemical classes of defensive compounds than cactus. Second, we compared the metagenomes of foregut contents and cecal contents in *N. stephensi* with the prediction that the abundance of detoxification genes would be higher in the foregut to facilitate detoxification prior to the major site of absorption in the small intestine.

### Comparative metagenomics

Total DNA was isolated from foregut and cecal contents using MoBio PowerFecal DNA extraction kit (MoBio, Carlsbad, California). Extracted DNA was sent to Argonne National Laboratory for sequencing. Genomic DNA was sheared using a Covaris Sonicator, to roughly 150 bp and metagenomic shotgun libraries were prepared using the Illumina TruSeq DNA sequencing preparation kits. Libraries were sequenced on the Illumina HiSeq2000 platform using a 2 × 100 bp run and V3 chemistry, which resulted in overlapping sequences.

Metagenomic sequences were uploaded to the MG-RAST website [23]. Sequences were screened against the genome of *Arabidopsis thaliana* to remove potential contamination from the plant-based diet. The reads were then filtered using dynamic trimming with a quality threshold of 15, such that any sequences with more than 5 low-quality bases were removed. Several foregut metagenomes contained high abundances of sequences (~30-50%) identified as plant DNA, likely resulting from the diet. Therefore, we partitioned our analysis by domain by selecting sequences identified as bacteria using the non-redundant multi-source protein annotation database, M5nr [24]. Abundances of non-plant eukaryotic and archaeal sequences were too low for sufficient analysis. Sequences identified as bacterial in origin were then annotated to identify putative functions using the KEGG Orthology database [25, 26] with the following thresholds: (1) e-values < le-5, (2) a minimum percent identity to database sequences of 60% and (3) a minimum alignment length of 15 bases. We compared the abundances at Level 3 of the KEGG hierarchical classification, as well as specific genes associated with “Xenobiotic degradation and metabolism”, “Geraniol degradation”, and “Limonene and pinene degradation”. Abundances of functional categories were compared using JMP 12.0 (SAS Institute Inc.) using t-tests and the robust Response Screening function that performs the Benjamini-Hochberg false discovery rate (FDR) correction for multiple tests [27].

We also conducted specific comparisons of the abundances of several detoxification genes: oxalyl-CoA decarboxylase, aryl alcohol dehydrogenase, and several genes associated with terpene metabolism. The gene oxalyl-CoA decarboxylase (*oxc*) degrades the compound oxalate. The gene aryl alcohol dehydrogenase is upregulated in the woodrat foregut when individuals are fed diets containing phenolic-rich resin from creosote bush [2]. Last, a number of genes associated with the “Limonene and pinene degradation” are enriched or present in the gut microbiota of specialist insect and avian herbivores that feed on terpene-rich plants [7, 20]. We compared the abundances of these genes using JMP 12.0 (SAS Institute Inc.) using pairwise t-tests (foregut *N. stephensi* vs. foregut *N. albigula*, and *N. stephensi* foregut vs. cecum) with the Dunn’s correction.

### Metagenome prediction

We also conducted microbial inventories by sequencing the variable regions 1-3 of 16S rRNA gene following experimental and bioinformatic methods described in detail elsewhere [28]. Taxonomic comparison of the communities between foregut and cecal chambers has already been conducted with larger sample sizes [11]. In the current study, we used 16S rRNA inventories to predict the metagenomic content of gut samples using genomic inference via PICRUSt [29]. We compared the relative abundances of functional categories between the shotgun metagenomic and predicted metagenomics to assess if predictive profiling via PICRUSt is informative for woodrat gut samples. First, we averaged the relative abundances of functional categories within a sample type (foregut and cecum), and transformed the relative abundances using a log(x+1) transformation. We then conducted a linear regression between transformed PICRUSt abundances and metagenomic abundances. We also plotted the relative abundances of functional categories that exhibited the highest fold difference between the foregut and cecal chambers to qualitatively inspect whether trends were similar for PICRUSt and metagenomic data.

### Accession Numbers

Metagenomic sequences can be found in the NCBI Short Read Archive (SRA) database under BioProject PRJNA336354, and on MG-RAST under Metagenome Project IDs mgp10174 and mgp10296. 16S rRNA sequences from the foregut and cecal chambers of *N. stephensi* that were used in PICRUSt analyses can be found in the SRA database under BioProject PRJNA449358.

## RESULTS

### Concentrations of PSCs across the gut

Water concentration varied over the length of the gut, with a significant reduction in the large intestine (Fig. 2). Oxalate concentrations in the entire gut were significantly less than in the food material (Fig 2; P < 0.0001). Oxalate concentrations also varied significantly across gut regions (P < 0.001), and were highest in the foregut and lowest in the stomach. Conversely, foregut concentrations of monoterpenes were not significantly lower than the food material (Fig. 2; P > 0.05 for all). Concentrations of monoterpenes did vary across gut regions, with the lowest concentrations always being in the cecum (Fig. 2; P < 0.0001 for all). In fact, camphene and β-pinene were not detectable in any cecal samples.

**Fig. 1.**
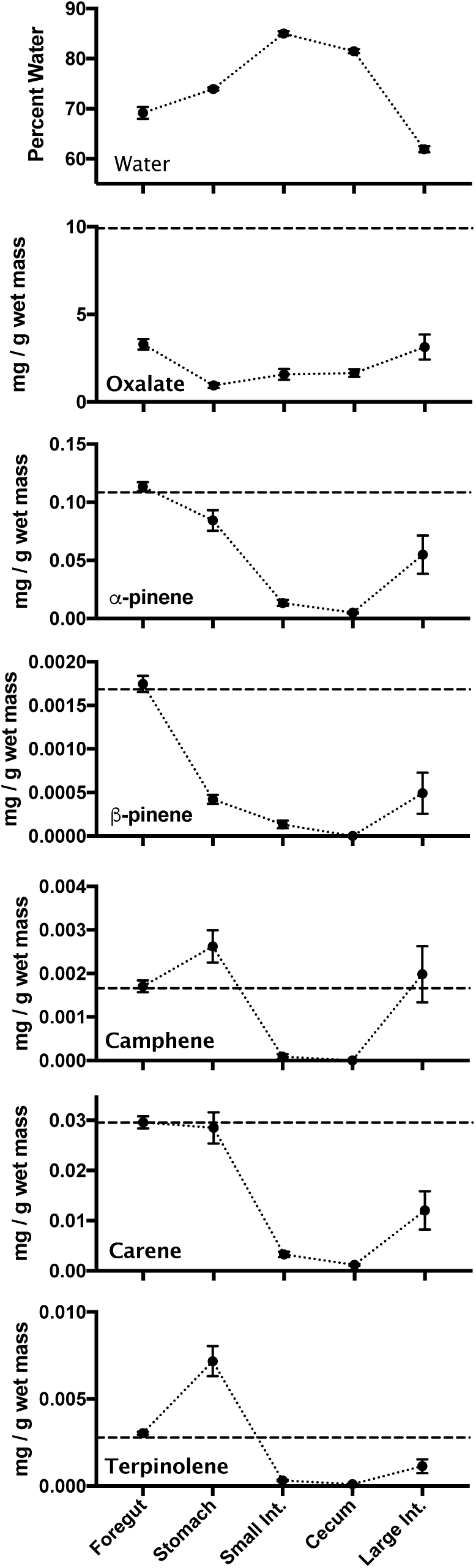
Concentrations of water and toxins across gut regions. The dotted line on toxin graphs represents concentrations as measured in the food material presented to woodrats. Points represent means ± 1 s.e.m.

**Fig. 2.**
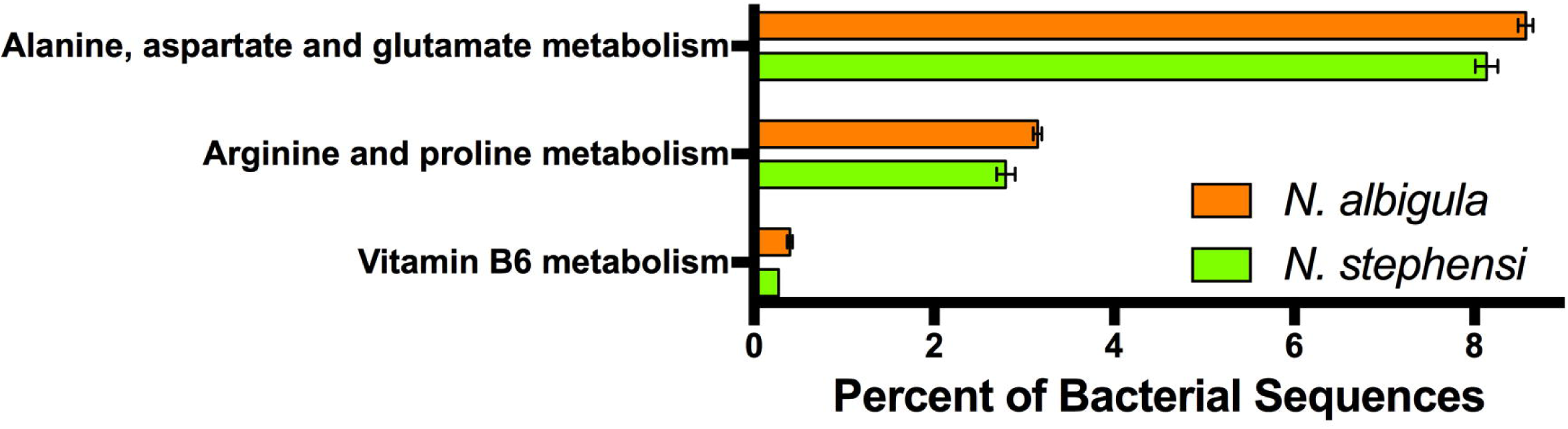
Functional categories (KEGG Level 3) that differed significantly between the foregut metagenomes of *N. albigula* and *N. stephensi*. Bars represent means ± 1 s.e.m.

### Metagenomic sequencing

We obtained an average of 3.35 million reads per sample (± 517 thousand). We first compared the bacterial metagenomes of foregut chambers from *N. stephensi* (which specializes on juniper) and *N. albigula* (which specializes on cactus). There were no functions that were significantly enriched in the foregut bacterial metagenomes of *N. stephensi*. The foregut bacterial communities of *N. albigula* were enriched in functions related to the metabolism of several amino acids and vitamin B6 (Fig. 3). Further, when investigating detoxification genes, there were no detectable differences between the two species.

**Fig 3.**
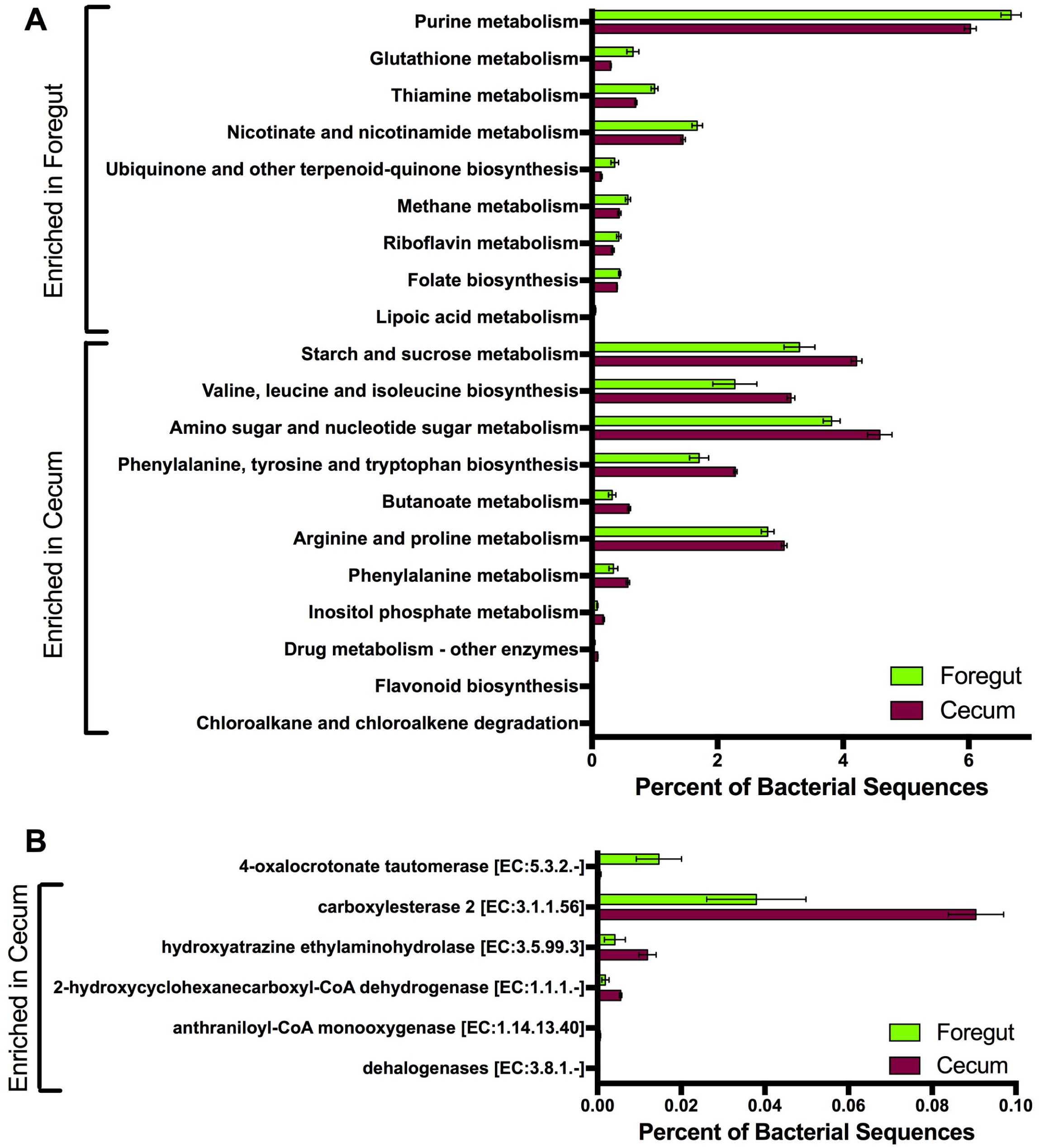
Differences in the metagenomic content of the foregut and cecal chambers of *N. stephensi*. Bars represent means ± 1 s.e.m. A) Functional categories (KEGG Level 3) that differed significantly in abundance between the foregut and cecal metagenomes. B) Detoxification genes that differed significantly in abundance between the foregut and cecal metagenomes.

We next searched for differential functions between the foregut and cecal chambers of *N. stephensi*. The foregut bacterial metagenome was enriched in genes associated with the metabolism of nucleotides (purines), amino acids (glutathione, thiamine), vitamins (riboflavin, nicotinate, folate), and lipoic acid (Fig. 4A). The cecal bacterial metagenome was enriched for genes associated with the metabolism of starch and sucrose, as well as the biosynthesis of several essential amino acids (Fig. 4A). The cecum was also enriched in several functional categories related to detoxification (drug metabolism, chloroalkane and chloroalkene degradation; Fig. 4A).

**Fig. 4.**
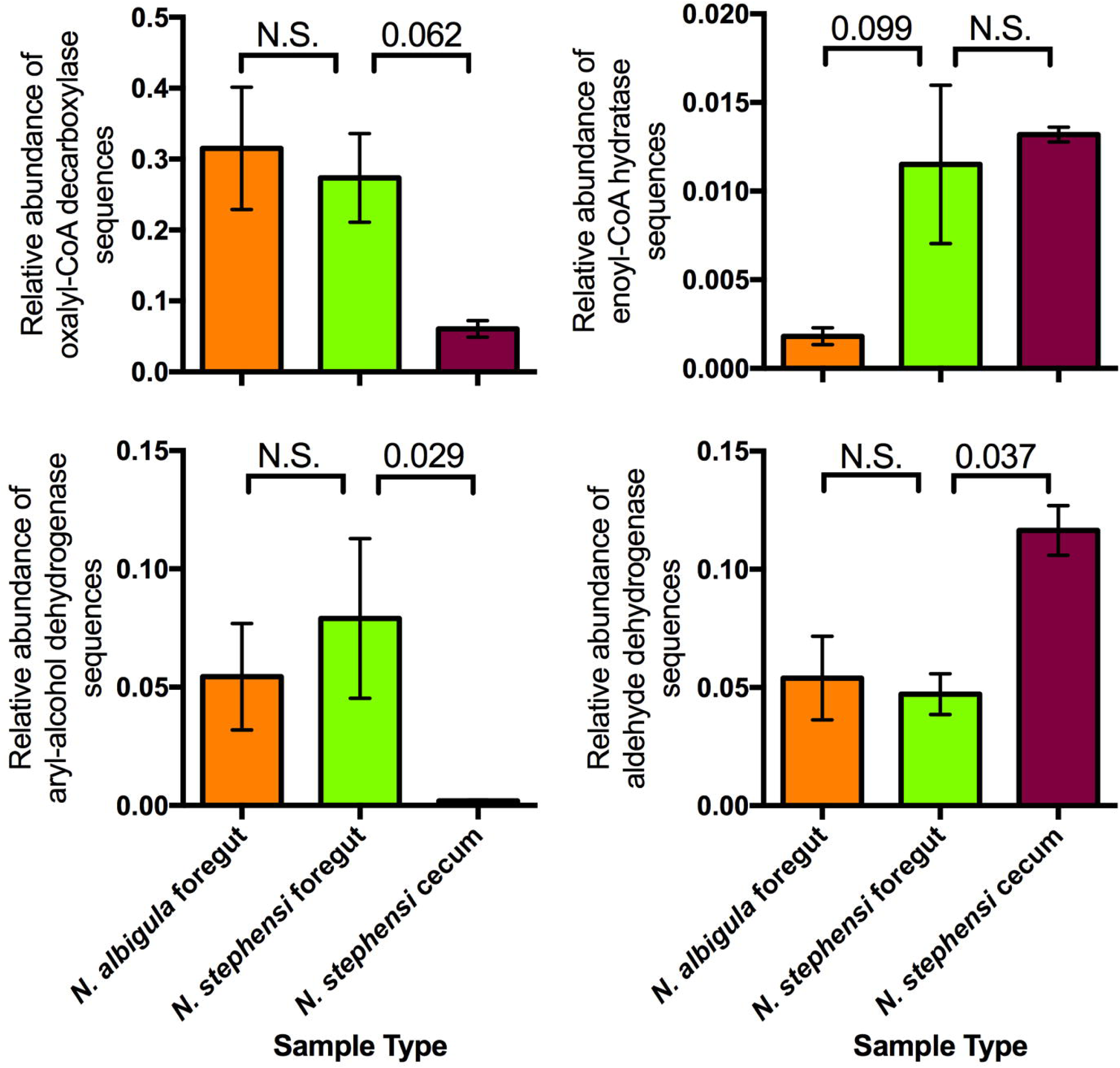
Relative abundances of specific detoxification enzyme genes associated with *a priori* hypotheses from other systems (see Methods). Bars represent means ± 1 s.e.m. Relative abundances were compared using t-tests (foregut *N. stephensi* vs. foregut *N. albigula*, and *N. stephensi* foregut vs. cecum) with the Dunn’s correction.

We also specifically investigated genes associated with “Xenobiotic degradation and metabolism”, “Geraniol degradation”, and “Limonene and pinene degradation”. Here, the only gene enriched in the foregut was 4-oxalocrotonate tautomerase, which was 24× more abundant in the foregut compared to the cecum (Fig. 4B). There were 5 other detoxification genes that were significantly more abundant in the cecum compared to the foregut chamber (Fig. 4B). Three of these genes (carboxylesterase 2, hydroxyatrazine ethylaminohydrolase, and 2-hydroxycyclohexanecarboxyl-CoA dehydrogenase) were 2-3 × more abundant in the cecal chambers compared to the foregut chambers. Two genes, anthraniloyl-CoA monooxygenase and dehalogenases, were not detected in the foregut chambers, but were present in the cecal chambers of the juniper-feeding *N. stephensi* (Fig. 4B).

Last, we focused on a subset of detoxification enzymes highlighted from previous studies. The gene oxalyl-CoA decarboxylase, which is anticipated to function in oxalate degradation, did not exhibit differential abundance between the foregut chambers of *N. stephensi* and *N. albigula*. However, the abundance of this gene was 4.5× higher in the foregut of *N. stephensi* compared to the cecum (Fig. 5). Similarly, aryl-alcohol dehydrogenase, which is more abundant in the woodrat foregut when animals are fed phenolic-rich creosote resin [2], did not exhibit significant differences between the foregut chambers of *N. stephensi* or *N. albigula*, but was 41 × higher in the *N. stephensi* foregut compared to the cecum (Fig. 5). Last, we investigated several genes in the “Limonene and pinene degradation” pathway that are enriched in other herbivores that feed on terpene-rich plants [7, 20, 21]. Enoyl-CoA hydratase (E.C. 4.2.1.17) was more abundant in the foregut of *N. stephensi* compared to *N. albigula*, with no difference between the foregut and cecal chambers of *N. stephensi* (Fig. 5). Additionally, aldehyde dehydrogenase (E.C. 1.2.1.3) exhibited no difference between the foregut chambers of *N. stephensi* and *N. albigula*, but was 2.5× more abundant in the cecal chambers of *N. stephensi* (Fig. 5).

**Fig. 5.**
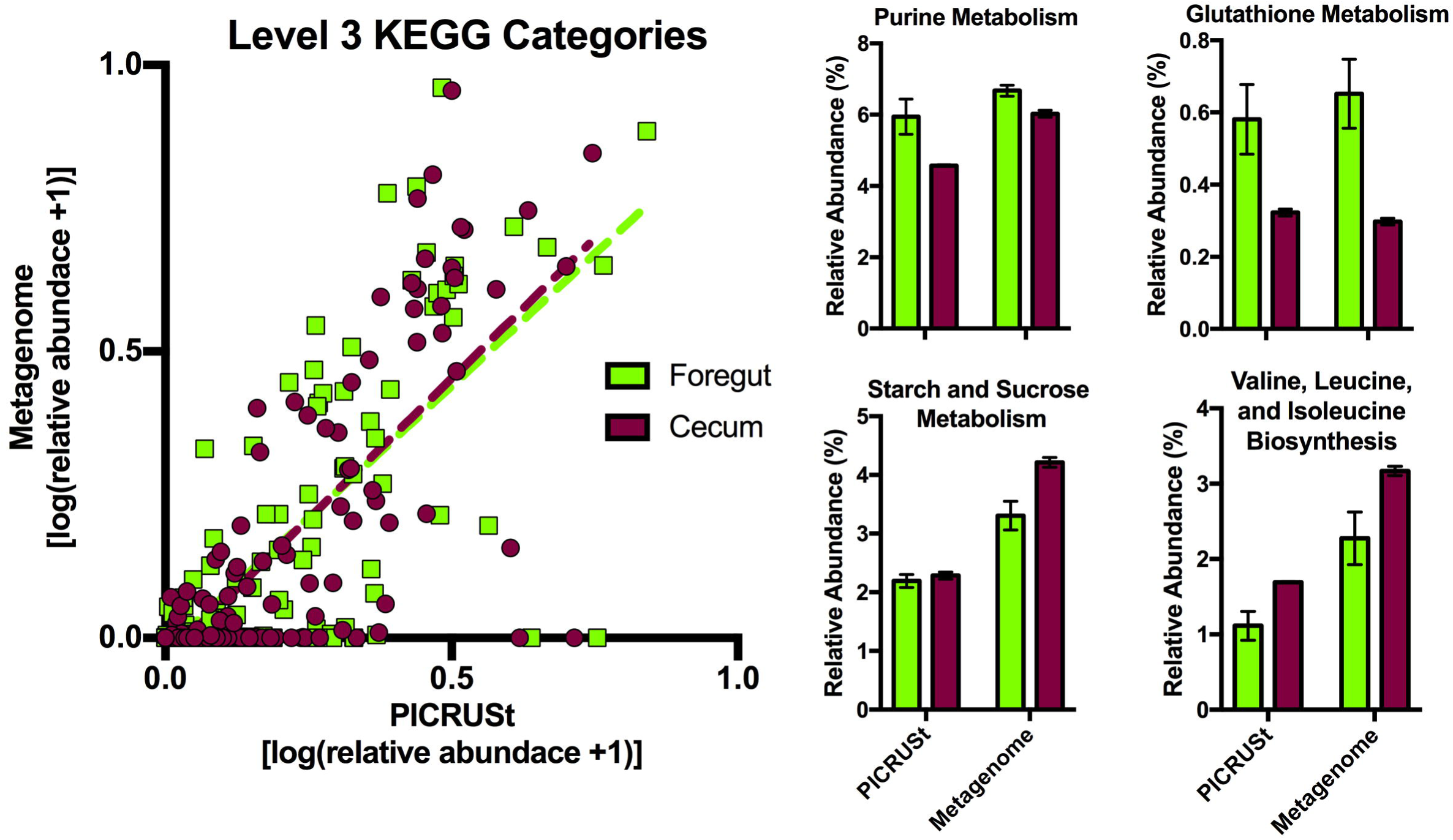
**A)** A comparison of predictive profiling by PICRUSt and metagenomic data. **B)** Bar graphs depict the functional categories that exhibited the highest fold-differences between the foregut and cecal metagenomes. Bars represent means ± 1 s.e.m.

We found a significant positive relationship between the relative abundances of functional categories as estimated by PICRUSt and determined by metagenomic sequencing (Fig. 6; foregut: R^2^ = 0.49, slope = 0.922 ± 0.074, P < 0.0001; cecum: R^2^ = 0.51, slope = 0.978 ± 0.072, P < 0.0001). Some functional categories, such as those representing ‘Peptidases’ and ‘Amino acid Related Enzymes’, were not well predicted by PICRUSt, in that they were present on the x-axis (predicted to be present by PICRUSt but not detected by metagenomic sequencing). However, categories that exhibited the greatest fold-differences between the two chambers were also predicted to differ by PICRUSt (Fig. 6; with the exception of Starch and Sucrose Metabolism).

## DISCUSSION

Here, we investigated the nature and location of microbial detoxification in the guts of Stephen’s woodrat, a juniper specialist. The location of detoxification within the gut seems to vary depending on the chemical class of defensive compounds. We discuss our findings below, as well as the potential ecological and evolutionary consequences of foregut vs hindgut detoxification by the microbiome.

We first compared concentrations of PSCs along the length of the gastrointestinal tract. We predicted that if toxins were metabolized early in the gut, their concentrations in the foregut chamber would be lower than those measured in food material, similar to what has been observed for usnic acid in the reindeer rumen [5]. This was observed in the case of oxalate, where foregut concentrations were roughly a third of those measured in the diet. Since oxalate is detected only at very low levels in the urine of woodrats [16] and mammals do not produce the necessary oxalate-degrading enzymes [30], it seems that oxalate is detoxified efficiently by microbes in the foregut. Conversely, terpene concentrations in the foregut were the same as those measured in food material, suggesting that the foregut is not a significant site for detoxification. Instead, concentrations of terpenes were lowest in the cecum, where some terpene compounds were not even detected. These results are similar to those observed in sage-grouse, where terpene concentrations are also lowest in the cecal chambers [31]. *Neotoma stephensi* has physiological adaptations that reduce terpene absorption in the gut [32], perhaps allowing terpene metabolism to be delayed until the food reaches the cecum. In the case of terpenes, concentrations increased in the large intestine compared to the cecum. While this change seems puzzling, we hypothesize that the selective absorption of water in this gut region results in higher concentrations of toxins. Further experiments, perhaps using radiolabeled compounds, are needed to understand the routes of absorption and detoxification of these compounds in different regions of the woodrat digestive system.

Interestingly, we did not observe many differences related to detoxification between the foregut microbial metagenomes of *N. stephensi* and *N. albigula*, even though these two host species feed on different diets with contrasting chemical profiles. It could be that the foregut microbiome offers broad detoxification capacities, while cecal microbial communities are more specialized. That being said, we did observe a higher abundance of enoyl-CoA hydratase in the foregut of *N. stephensi* compared to *N. albigula*. This gene plays a role in the ability for microbes to degrade terpenes [33], and is enriched in the gut metagenomes of sage-grouse [7] and mountain pine beetles [20]. Thus, there may be some functional enrichment for detoxification in the foregut of *N. stephensi*.

Within *N. stephensi*, the foregut and cecal microbial communities appear to provide different detoxification functions. In the foregut, we observed higher abundance of the gene 4-oxalocrotonate tautomerase, which is involved in the degradation of aromatic hydrocarbons by bacteria [34]. The foregut also had higher abundances of the gene oxalyl-CoA decarboxylase, which degrades oxalate, and aryl-alcohol dehydrogenase, which degrades aromatic hydrocarbons. The abundance of aryl-alcohol dehydrogenase increases in the foregut of another woodrat species (*N. lepida*) when animals are fed diets containing phenolic PSCs [2] and thus may be an important enzyme that is provided by the foregut microbial community.

We also obtained evidence for enrichment of detoxification enzymes in the cecal chamber. Higher-level gene categories, such as ‘Drug Metabolism’ and ‘Chloroalkane and Chloroalkene Degradation’ were more abundant in the cecal chambers than the foregut. Specifically, the cecum was enriched for genes such as hydroxyatrazine ethylaminohydrolase, which deaminates or dechlorinates triazine compounds [35, 36], and anthraniloyl-CoA monooxygenase, which plays a role in the catabolism of aromatic compounds [37]. Additionally, the aldehyde dehydrogenase gene was significantly more abundant in the cecum. This gene is annotated as a component of the ‘Limonene and Pinene Degradation’ pathway, and is found to be enriched in the gut metagenomes of sage-grouse [7] and mountain pine beetles [20].

Finally, we tested whether predictive profiling of metagenomics based on 16S rRNA sequences is appropriate for woodrat gut samples. We found a significant relationship between the relative abundances of functional categories as estimated by PICRUSt and determined by metagenomic sequencing. Additionally, results for PICRUSt and metagenomics were consistent for some of the categories that exhibited the largest differences between the two chambers. However, PICRUSt did not predict all of the categories accurately. Many functional categories (e.g. ‘Peptidases’ and “Amino acid Related Enzymes’) were predicted to be present by PICRUSt analysis, but were not detected by metagenomic sequencing. Interestingly, the gut microbial communities of herbivores tend to have lower abundances of genes encoding protease relative to carnivores, likely to conserve these nutrients on a low protein diet [38]. Thus, the use of predictive metagenomic profiling may not always be feasible for non-model systems, especially given that most microbial genomes used for predictive profiling are from the human gut. Indeed, PICRUSt works significantly better for human samples than for samples from nonhuman mammalian guts [29]. The sequencing of additional genomes from nonhuman mammals would help to improve the accuracy of programs like PICRUSt in non-model systems. It seems that results from predicting profiling in non-model systems should perhaps be verified with other methods, such as quantitative PCR.

Overall, we hypothesized that microbial detoxification should occur proximally in the gastrointestinal tract, so as to facilitate degradation of PSCs before they would be absorbed later in the gut. However, it seems that detoxification processes may be spatially structured along the length of the gut. The evidence for foregut detoxification is strongest for oxalate, where abundances of the detoxification enzyme (oxalyl-CoA decarboxylase) were high, and concentrations of oxalate were already much lower than in food material, and abundances of potentially oxalate-degrading bacteria are highest [19]. Terpene degradation may occur in the cecal chamber, where abundances of terpene-degradation genes were higher, and concentrations of terpenes were at their lowest. It is somewhat puzzling why this spatial arrangement would exist, given that terpenes are absorbed very rapidly due to their small size and high lipophilicity [32]. However, *N. stephensi* exhibits lower terpene absorption than other woodrat species [32], resulting in relatively higher terpene exposure and resultant detoxification by microbial communities in the hindgut. Laboratory rats are able to excrete ~25% of absorbed monoterpenes back into the gut lumen through the biliary duct [39], an excretion route that may be enhanced in a specialist herbivore such as *N. stephensi*. This excretion route may facilitate further microbial degradation of terpenes in the hindgut. While we did not measure concentrations of phenolics through the gut, various genes associated with degrading aromatic hydrocarbons were enriched in both the foregut and cecum. Phenolics can bind to dietary protein and digestive enzymes, thus limiting nutrient availability [40]. Therefore, foregut detoxification of phenolics would be beneficial for avoiding the anti-nutritive effects of PSCs. Again, labeled chemicals to trace sites of absorption, excretion, and metabolism would enhance our understanding of the location of these processes.

The location of detoxification within the gut could have important ecological and evolutionary implications for plants and herbivores. For example, when comparing several species of rodents, only those with foregut chambers were able to metabolize significant amounts of oxalate [41]. Thus, evolution of this chamber may be important for the ability to consume oxalate-rich plants. Consumption of plant toxins might also drive changes in gut anatomy to enhance detoxification by facilitating a longer duration of residency in various gut chambers. For example, woodrats feeding on phenolic-rich oak exhibit an increase in cecal size [42], which could be driven by increased fiber content of oak or to facilitate the detoxification of PSCs. Interestingly, supplementing diets with isolated phenolic compounds causes an increase in cecal size in lab rats [43]. Alterations of the size or function of gut chambers in response to toxins could have implications for dietary niche breadth or energetics. In herbivorous birds, there is evidence suggesting that microbial detoxification occurs in both foregut [6] and hindgut chambers [7]. The guts of birds are under strong selective pressure to be small so as to facilitate flight [44], and thus sites of microbial detoxification may be especially important in these species. Studies on additional and phylogenetically diverse herbivores and classes of PSCs may help us to better understand the importance of microbial detoxification in the gut, and how the spatial arrangement of these processes contribute to animal morphology, performance, ecology, and evolution.

## Acknowledgements

We would like to thank Madelina James and KayLene Yamada for helping with feeding trials and Chelsea Merriman for analysis of monoterpenes. Funding for this study was provided by the National Science Foundation (Doctoral Dissertation Improvement Grant, DEB 1210094, to M.D.D. and K.D.K.; DEB 1342615 to M.D.D.; DEB-1146194 to JSF; and DBI 1400456 to K.D.K).

